# AICAR induces apoptosis and inhibits migration of prostate cancer cells through an AMPK/mTOR-dependent pathway

**DOI:** 10.1101/474197

**Authors:** Chia-Cheng Su, Kun-Lin Hsieh, Shu-Chi Wang, Hsin-Chih Yeh, Shu-Pin Huang, Po-Len Liu, Shih-Hua Fang, Wei-Chung Cheng, Kuan-Hua Huang, Fang-Yen Chiu, I-Ling Lin, Ming-Yii Huang, Chia-Yang Li

## Abstract

AICAR (5-aminoimidazole-4-carbox-amide-1-β-D-ribofuranoside), an AMP-activated protein kinase (AMPK) agonist, has demonstrated antitumor activities for several types of cancers. However, the activity of AICAR on the cell growth and metastasis of prostate cancer has not been extensively studied. Herein we examine the effects of AICAR on the cell growth and metastasis of prostate cancer cells, 22RV1 cells. Cell growth was performed by MTT assay and soft agar assay. Cell apoptosis was examined by Annexin V/PI staining and PARP cleavage Western blot. Cell migration was evaluated by wound-healing assay. The expression of EMT-related protein and the activity of the AMPK/ mTOR-dependent pathway were analyzed by Western blot. In addition, we also tested the effect of AICAR on the chemosensitivity to docetaxel using MTT assay. Our results indicated that AICAR inhibits cell growth, induces apoptosis, attenuates TGF-β-induced cell migration and EMT-related protein expression, and enhances the chemosensitivity to docetaxel through regulating the AMPK/mTOR-dependent pathway. Collectively, these findings support AICAR as a potential therapeutic agent for the treatment of prostate cancer.

## Introduction

Prostate cancer is the most common cancer and the second leading cause of cancer-related death among men in the United States [1]. Current treatment options for prostate cancer include surgery, hormonal therapy, chemotherapy, radiation therapy, radiofrequency ablation, high-intensity focused ultrasound, cryotherapy, and cancer vaccine [2, 3]. Androgen deprivation therapy (ADT) by surgical or chemical castration has been the mainstay of treatment for advanced prostate cancer in the past few decades. However, the majority of androgen-sensitive prostate cancer patients will eventually develop resistance to ADT within 1 to 3 years and the disease will become androgen-independent [4]. Currently, there is no effective therapy for recurrent prostate cancer. Hence, a novel therapeutic method for prostate cancer is needed.

The 5’-adenosine monophosphate (AMP)-activated protein kinase (AMPK) is activated by increases in cellular ATP/AMP ratio and plays an important role in regulating glycolytic activity and maintaining energy balance at both cellular and whole body levels [5]. Previous studies indicated that AMPK exhibits intricate relations with other energy/metabolite sensor pathways (e.g. SIRT1, Akt, mTOR, PARPs, etc.) and acts in a coordinated fashion with these [6-10]. AMPK induction leads to enhanced mitochondrial oxidation and mitochondrial biogenesis that have been shown to exert anti-Warburg and anti-proliferative effects in several types of cancers, such as leukemia [11], breast cancer [12], pancreatic cancer [13], hepatocellular carcinoma [14], and prostate cancer [15]. These findings suggest that AMPK activation may be used beneficially for cancer treatment.

The adenoside analog compound, 5-aminoimidazole-4-carbox-amide-1-β-D-ribofuranoside (AICAR), is intracellularly converted by adenosine kinase to the non-phosphorylated derivative amino-imidazolecarboxamide ribonucleotide (ZMP), an analogue of AMP, which activates AMPK [16]. Therefore, AICAR is often used as an activator of AMPK [17] to modulate cellular energy homeostasis, although AMPK-independent effects have also been proposed [18, 19]. A previous study indicated that AICAR inhibited the growth of androgen-independent (DU145, PC3) and androgen-sensitive (LNCaP) cells after four days of treatment [20]. In addition, AICAR inhibited two key enzymes involved in protein synthesis, mTOR and p70S6K, and blocked the ability of the androgen R1881 to increase cell growth and the expression of two enzymes for *de novo* fatty acid synthesis, acetyl CoA carboxylase and fatty acid synthase, in the LNCaP cells [20]. A current study showed that AICAR induced AMPK-independent programmed necrosis in prostate cancer cells [21]. Collectively, these studies indicate that AICAR has potential in inhibiting the growth of prostate cancer. However, the effect of AICAR on the apoptosis and migration of prostate cancer remains unclear. In the present study, we evaluated the effects of AICAR on apoptotic activity, migration and chemosensitivity through AMPK phosphorylation in human prostate cancer cells.

## Materials and Methods

### Reagents

RPMI 1640 medium, penicillin, streptomycin and fetal bovine serum (FBS) were purchased from Gibco-BRL (Life Technologies, Grand Island, NY, USA). AICAR, bovine serum albumin (BSA), phosphate-buffered saline (PBS), RIPA buffer, protease inhibitor cocktail, phosphatase inhibitor cocktail, stripping buffer, thioglycollate medium, and 3-(4,5-dimethylthiazol-2-yl)-2, 5-diphenyl tetrazolium bromide (MTT) were purchased from Sigma Aldrich (St. Louis, MO, USA). AICAR was purchased from Cayman Chemical (Ann Arbor, MI, USA). Alexa Fluor® 488 Annexin V/Dead Cell Apoptosis Kit was purchased from Thermo Fisher Scientific (Eugene, Oregon, OR, USA). BCA protein assay reagent was purchased from Thermo Scientific (Waltham, MA, USA). For western blotting, rabbit antibodies against human phospho-AMPK, AMPK, MYC, mTOR, PARP, phospho-p70S6K, p70S6K, TSC-1, TSC-2, β-actin and secondary antibodies were purchased from Cell Signaling (Farmingdale, NY, USA). Caspase-Glo 3/7 assay kit was purchased from Promega (Madison, WI, USA). Transforming growth factor-beta 1 was purchased from PeproTech (Rocky Hill, NJ, USA). Docetaxel (Taxothere, 20 mg/mL) was obtained from Sanofi Aventis (Germany).

### Cell culture

Human prostate cancer cell line, 22Rv1, purchased from Bioresource Collection and Research Center (Food Industry Research and Development Institute, Hsinchu, Taiwan), and was cultured in RPMI 1640 supplemented with antibiotics (100 U/mL penicillin and 100 U/mL streptomycin) and 10% (v/v) FBS in a humidified atmosphere of 5% CO_2_ at 37°C and passaged every 2-3 days to maintain growth.

### MTT assay

22Rv1 cells were seeded in a 96-well plate at a concentration of 1 × 10^5^ cells, and were allowed to acclimatize overnight. Cells were treated with various concentrations of the AICAR (0, 0.5, 1 and 3 mM) for 24 hr. Cell viability was measured by the ability of viable cells to reduce MTT to formazan based on the ability of living cells utilized thiazolyl blue and converted it into purple formazan. The concentration of formazan is measured by determining the OD at 570 nm using a microplate reader (BioTek Instruments, Inc., Winooski, VT, USA). The results are given as relative percentage to the untreated control. To detect the synergistic effects of AICAR, cells were treated with different doses of the AICAR (0, 0.5, and 1 mM) combined with different doses of docetaxel for 24 and 48 hr.

### Soft agar colony formation assay

Noble agar (BD Biosciences, Franklin Lakes, NJ, USA) was dissolved in complete medium and coated with 0.5% agar solution on the bottom of 6-well plates. After solidifying, top agar medium mixture (0.3%) containing 5 × 10^3^ cells was added, and incubated at 37°C in a humidified atmosphere of 5% CO_2_ for 3 weeks. Colonies were stained with 0.05% crystal violet-10% ethanol in PBS. Photographs of the stained colonies were captured using Bio-Rad ChemiDoc XRS^+^ system (Bio-Rad Laboratories, Inc., Hercules, CA, USA) and quantified using ImageJ software (National Institutes of Health, Bethesda, MD, USA).

### Apoptosis assay

22Rv1 cells were treated with different doses of AICAR (0, 0.5, 1, and 3 mM) for 24 hr. Cells were trypsinized, washed twice by cold PBS, and stained with Alexa Fluor® 488 Annexin V and propidium iodide (PI) according to manufacturer’s protocol (Thermo Fisher Scientific, Rockford, IL, USA). Apoptotic cells were determined using FC500 flow cytometer (Beckman-Coulter, Fullerton, CA, USA). Ten thousand events were collected per sample. Data were analyzed by CXP analysis software (Beckman-Coulter, Fullerton, CA, USA).

### Western blot

Cells were lysed by RIPA buffer with protease inhibitors and phosphatase inhibitors according to the manufacturer’s protocol (Sigma Aldrich, St. Louis, MO, USA). The protein concentration was determined using the BCA protein assay reagent according to the manufacturer’s instructions (Thermo Scientific, Waltham, MA, USA). Cellular protein extracts were separated by electrophoresis using 8% SDS polyacrylamide gel and were electroblotted onto PVDF membranes. The membranes were incubated with blocking solution for 1 hr at room temperature, followed by incubation overnight with primary antibodies at 4°C (1:1000). Blots were washed three times with Tris-buffered saline/Tween 20 (TBST) and incubated with a 1:5000 dilution of horseradish peroxidase conjugated secondary antibody for 1 hr at room temperature. Blots were again washed three times with TBST and developed using an ECL chemiluminescence substrate (Thermo Scientific, Waltham, MA, USA). The signals were captured and the band intensities were quantified using Bio-Rad ChemiDoc XRS^+^ system (Bio-Rad Laboratories, Inc., Hercules, CA, USA).

### Caspase 3/7 activity assay

22Rv1 cells were seeded in a 96-well white plate at a concentration of 2.5 × 10^4^ cells, and were allowed to acclimatize overnight. Cells were treated with various concentrations of the AICAR (0, 0.5, 1, and 3 mM) for 24 hr. A total of 100 μL Caspase-Glo® 3/7 reagent was added to each well, gently mixed contents of wells using a plate shaker at 300~500 rpm for 30 seconds, and then samples were incubated for 30 min at RT. Enzyme activity is directly proportional to luminescence, which was measured using a luminescence microplate reader (BioTek Instruments, Inc., Winooski, VT, USA). Data were normalized relative to the caspase 3/7 activity of cells treated with DMSO alone.

### Migration assay

SPLScar^TM^ Block (SPL life sciences, Korea) was placed in a 24-well plate, seeded 5 × 10^4^ cells in each side of block, and was allowed to acclimatize overnight. The block was removed, replaced flash medium, and treated cells with 5 ng/ml TGF-β1 and different concentrations of AICAR. Cell migration was monitored under a phase-contrast microscope and the migratory distance was calculated.

### Statistical analysis

All data are expressed as means ± SD. Each value is the mean of three independent experiments. Statistical analysis was assessed via Student’s *t*-test using IBM SPSS Statistics v.19 (IBM Corp., Armonk, NY, USA), and the significant difference was set at \**p* < 0.05; \*\**p* < 0.01; \*\*\**p* < 0.001.

## Results

### AICAR inhibits the growth of prostate cancer cells

The 22Rv1 cell line is an androgen-independent prostate cancer epithelial cell line that is representative of clinical recurrent prostate cancer [22]. Firstly, we examined the effect of AICAR on the growth of 22Rv1 prostate cancer cells. Cells were treated with various concentrations (0, 0.5, 1 and 3 mM) of AICAR for 24 hr. Cell viability was analyzed by MTT assay. The experimental results showed that AICAR inhibits the growth of 22Rv1 cells in a dose-dependent manner (Fig. 1A). In addition, we further examined whether AICAR affects the colony growth under anchorage-independent condition. Cells were treated with different concentrations (0, 0.25, 0.5 and 1 mM) of AICAR and then conducted a soft agar assay. The experimental results showed that AICAR suppressed growth of 22Rv1 cells in soft agar in a dose-dependent manner (Fig. 1B).

**Figure 1.**
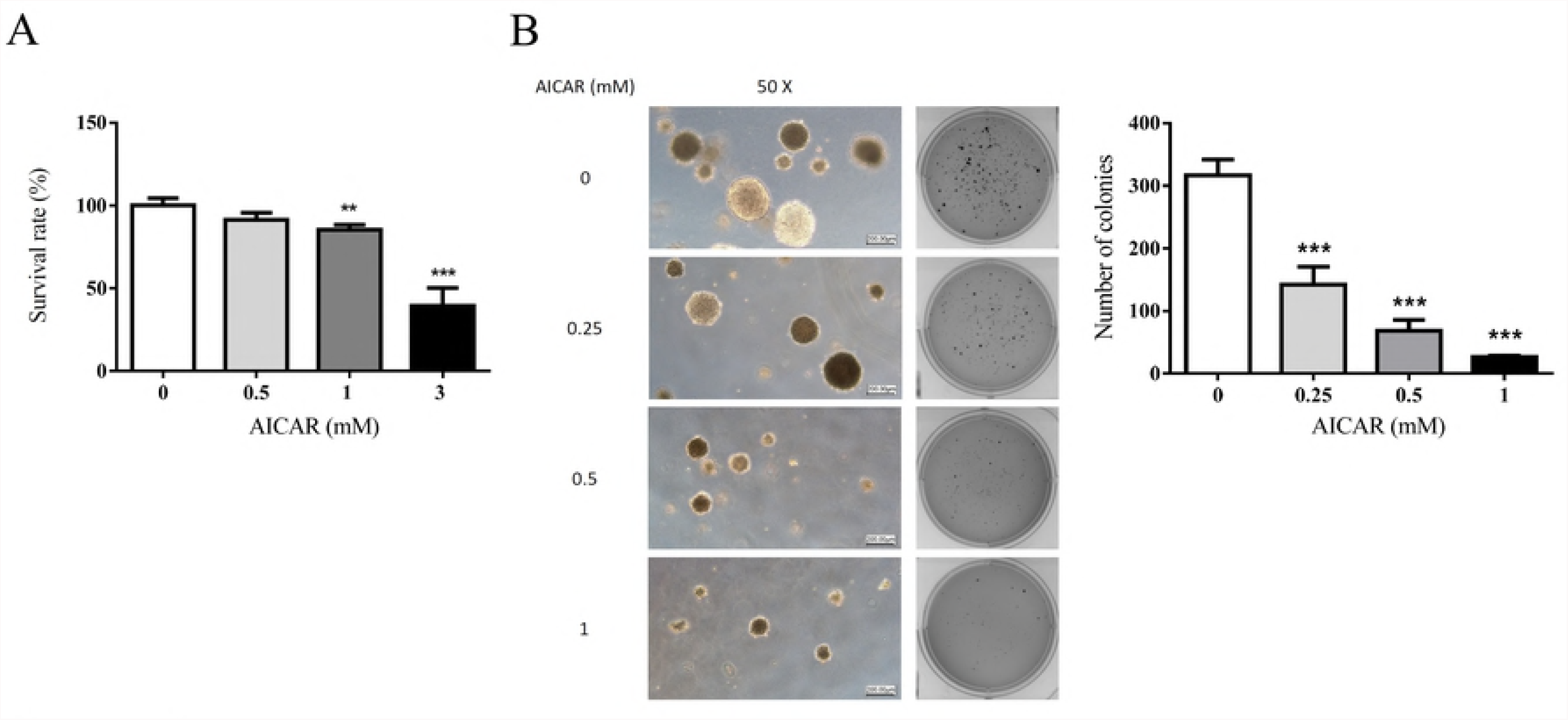
The effect of AICAR on the growth of prostate cancer cells. (A) 22Rv1 cells were treated with different concentration of AICAR for 24 hr. The cell viability was measured by the MTT assay. (B) 22Rv1 cells were treated with different concentration of AICAR, and then grown in soft agar, an anchorage-independent condition, for 3 weeks. Colonies were stained with crystal violet and captured using Bio-Rad ChemiDoc XRS^+^ system. Data are represented as means ± SD of triplicate values and statistical significance was determined using the Student’s *t*-test (\*\**p*< 0.01; \*\*\**p*< 0.001).

### AICAR induces apoptosis in prostate cancer cells

To further test whether AICAR induces apoptosis in prostate cancer cells, 22Rv1 cells were treated with various concentrations (0, 0.5, 1 and 3 mM) of AICAR for 24 hr. Apoptotic cells were detected with Annexin V/PI staining using flow cytometry. Our results demonstrated that AICAR induced apoptosis (Fig. 2A). Poly(ADP-ribose) polymerase (PARP) is a nuclear enzyme involved in DNA repair which is cleaved by caspase-3 during apoptosis [23]. We further examined whether AICAR affects the expression of PARP in 22Rv1 cells. As shown in Figure 2B, AICAR increased the expression of cleaved PARP, an apoptosis marker, in 22Rv1 cells. In addition, we also examined the activity of caspase 3/7 using a luminescent substrate based assay. Our results indicated that AICAR increased the activity of caspase 3/7 in 22Rv1 cells (Fig. 2C).

**Figure 2.**
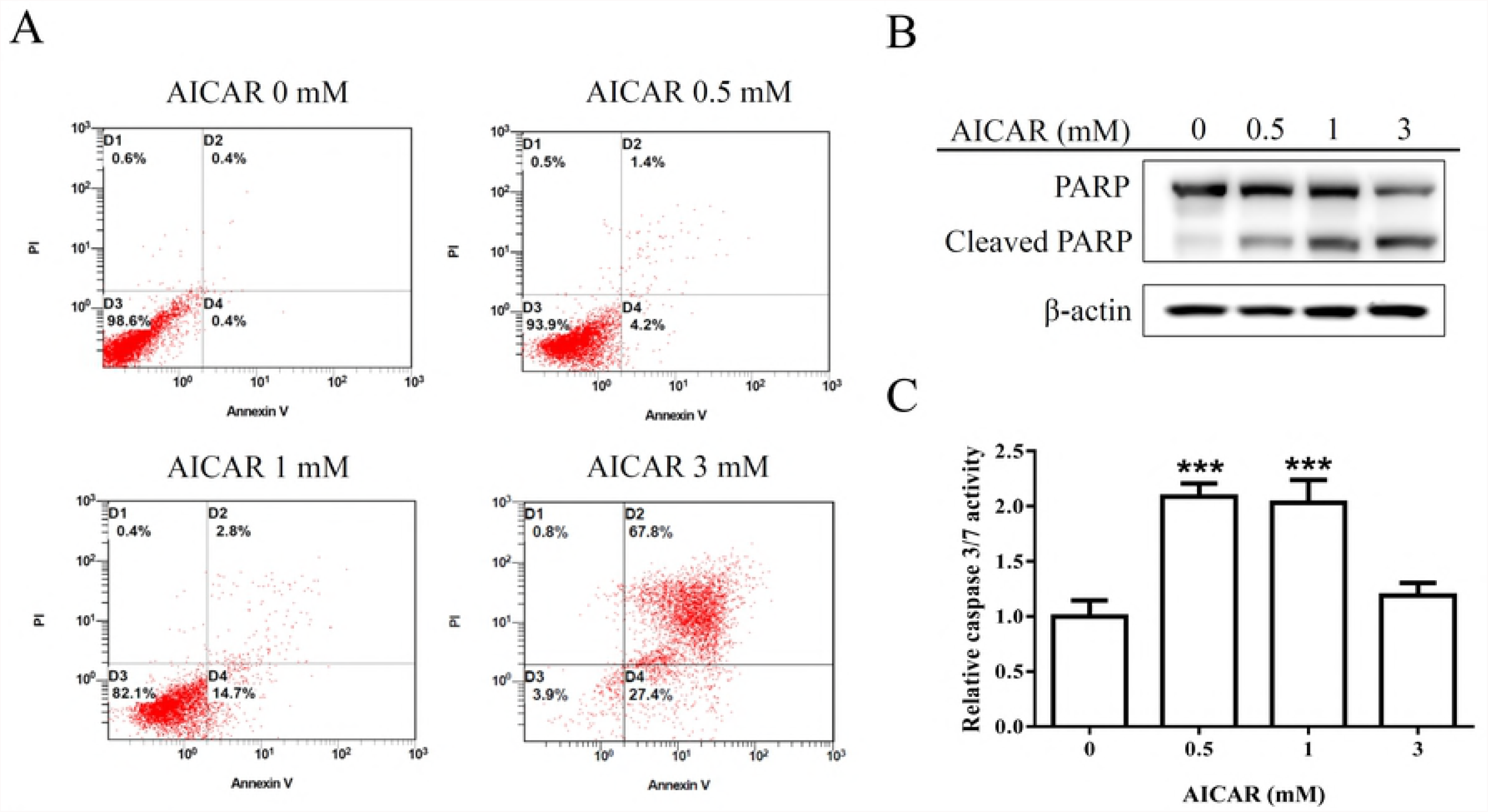
Effect of AICAR on the apoptosis in 22Rv1 prostate cancer cells. Cells were incubated with different concentrations of AICAR for 24 hr. (A) Cells were collected, stained with Annexin V and PI, and analyzed by flow cytometry. Data are representative of at least three independent experiments with similar results. (B) The expression of PARP was determined by Western blot. Actin was used as a loading control in Western blot. (C) Cellular caspase 3/7 activities were analyzed with caspase-glo assay kit. Data are represented as means ± SD of triplicate values and statistical significance was determined using the Student’s *t*-test (\*\*\**p*< 0.001).

### AICAR inhibits transforming growth factor-beta (TGF-β)-induced migration and epithelial to mesenchymal transition (EMT)

TGF-β signaling is well known as a key regulator of many biological processes in prostate cancer including inducing EMT, migration and metastasis [24]. To examine whether AICAR affects TGF-β-induced migration and EMT in prostate cancer cells, 22Rv1 cells were treated with 5 ng/mL TGF-β and various concentrations (0, 0.25 and

0.5 mM) of AICAR. Cell migration was performed by counting the average wound space. Our results showed that AICAR significantly inhibited TGF-β-induced migration (Fig. 3A and 3B). In addition, we also analyzed the expression of EMT-related proteins using Western blot. As shown in Figure 3C, AICAR inhibited TGF-β-induced EMT through inhibiting the expression of mesenchymal marker, N-cadherin, and enhancing the expression of epithelial marker, E-cadherin.

**Figure 3.**
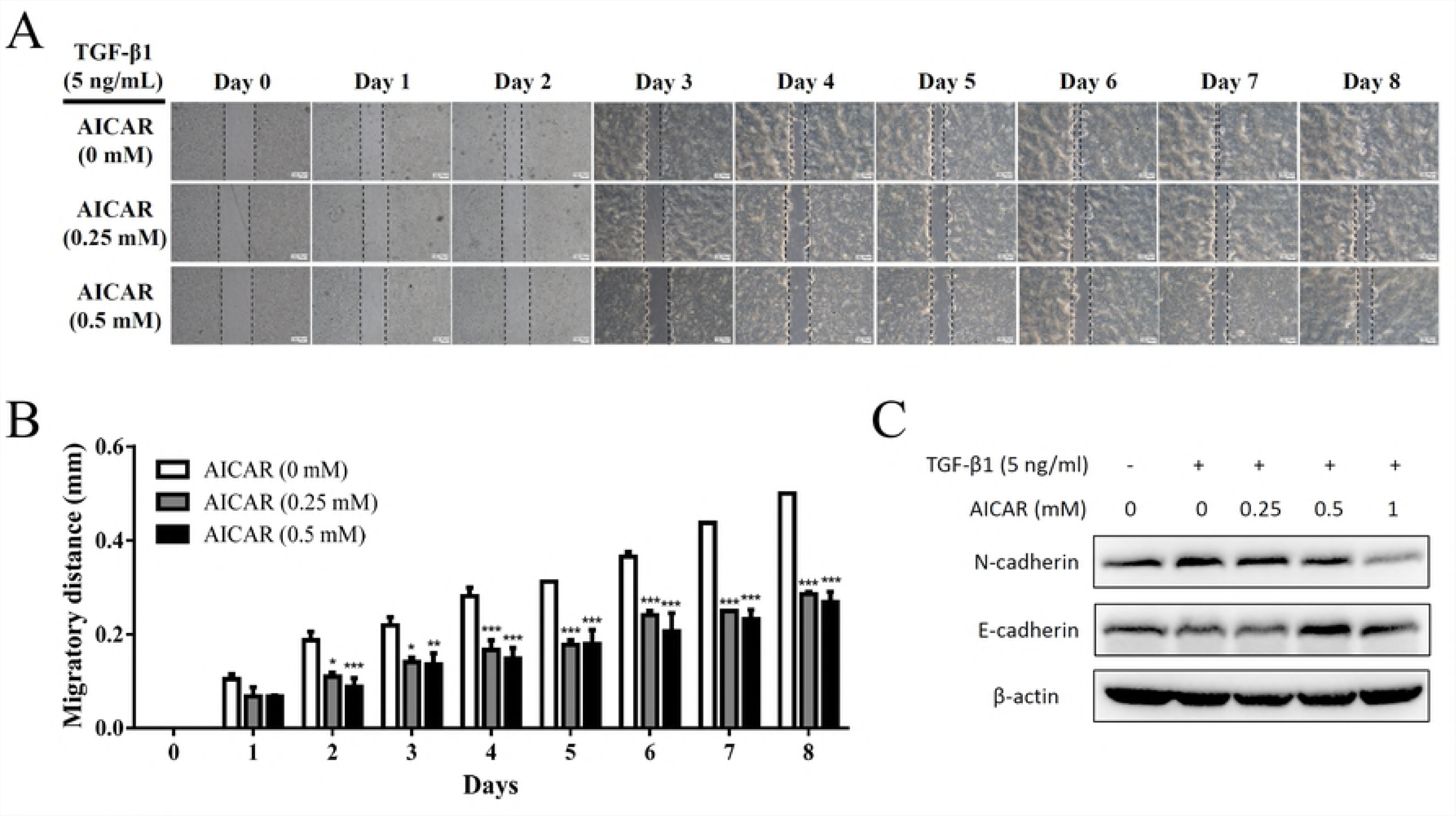
Effect of AICAR on the cell migration activity in 22Rv1 prostate cancer cells. Cells were seeded in SPLScar^TM^ Block overnight, the block was then removed and cells were treated with 5 ng/ml TGF-β1 and different concentrations of AICAR. (A) Cell migration was monitored under a phase-contrast microscope. Data are representative of at least three independent experiments with similar results. (B) The migratory distance was calculated. Data are represented as means ± SD of triplicate values and statistical significance was determined using the Student’s *t*-test (\**p*< 0.05; \*\**p*< 0. 01; \*\*\**p*< 0.001). (C) The expression of N-cadherin and E-cadherin was analyzed by Western blot. Actin was used as a loading control in Western blot. The Western blotting results are representative of results obtained in three separate experiments.

### AICAR exhibits synergistic effect with docetaxel treatment

Docetaxel-based therapy has been applied as the first-line chemotherapy for metastatic castration-resistant prostate cancer (mCRPC) for a decade due to its ability to improve the median overall survival (OS) by around 3 months [25, 26]. To examine whether AICAR affects the therapeutic efficiency of docetaxel-based therapy in prostate cancer, 22Rv1 cells were treated with various doses (0, 0.5 and 1 mM) of AICAR in the presence or absence of docetaxel for 24 hr. Cell survival was analyzed by MTT assay. As shown in Fig. 4, AICAR exhibited synergistic effect with docetaxel treatment in prostate cancer cells.

**Figure 4.**
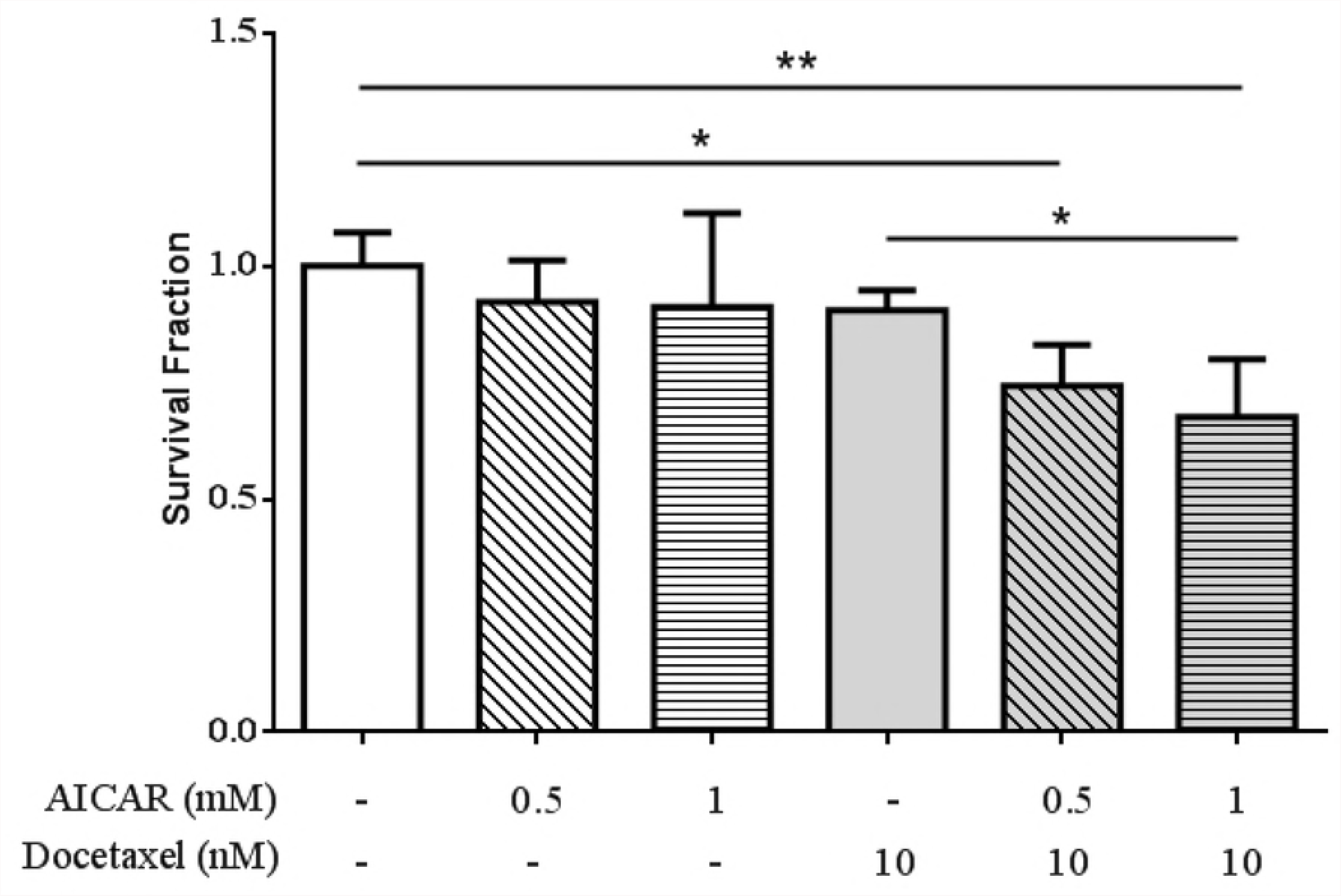
Effect of AICAR on the sensitivity of 22Rv1 prostate cancer cells to docetaxel treatment. Cells were treated with different concentrations of AICAR in the presence or absence of docetaxel for 24 hr. The cell viability was measured by the MTT assay. Data are represented as means ± SD of triplicate values and statistical significance was determined using the Student’s t -test (\**p*< 0.05; \*\**p*< 0.01).

### AICAR inhibits the growth of prostate cancer cells through activating an AMPK/mTOR-dependent pathway

AMPK is an upstream regulator in modulating the activation of TSC1 and TSC2 which consequently inhibits cell survival and proliferation via repressed mTOR-p70S6K-MYC signaling pathway [27, 28]. To investigate the mechanism of action of AICAR in regulating the activity of AMPK-dependent pathway, 22Rv1 cells were treated with different concentrations (0, 0.5, 1 and 3 mM) of AICAR. The expression of phospho-AMPK, AMPK, TSC-1, TSC-2, mTOR, phospho-p70S6K, p70S6K and MYC was analyzed by Western blot. As shown in Fig. 5A, AICAR promoted the phosphorylation of AMPK in 22Rv1 cells. In addition, our experimental results showed that AICAR enhanced the expression of TSC1 and TSC2 (Fig. 5B), whereas AICAR reduced the expression of mTOR and MYC as well as decreased the phosphorylation of p70S6K (Fig. 5C). These results suggest that AICAR inhibits the growth of prostate cancer cells through an AMPK/mTOR-dependent pathway (Fig. 5D).

**Figure 5.**
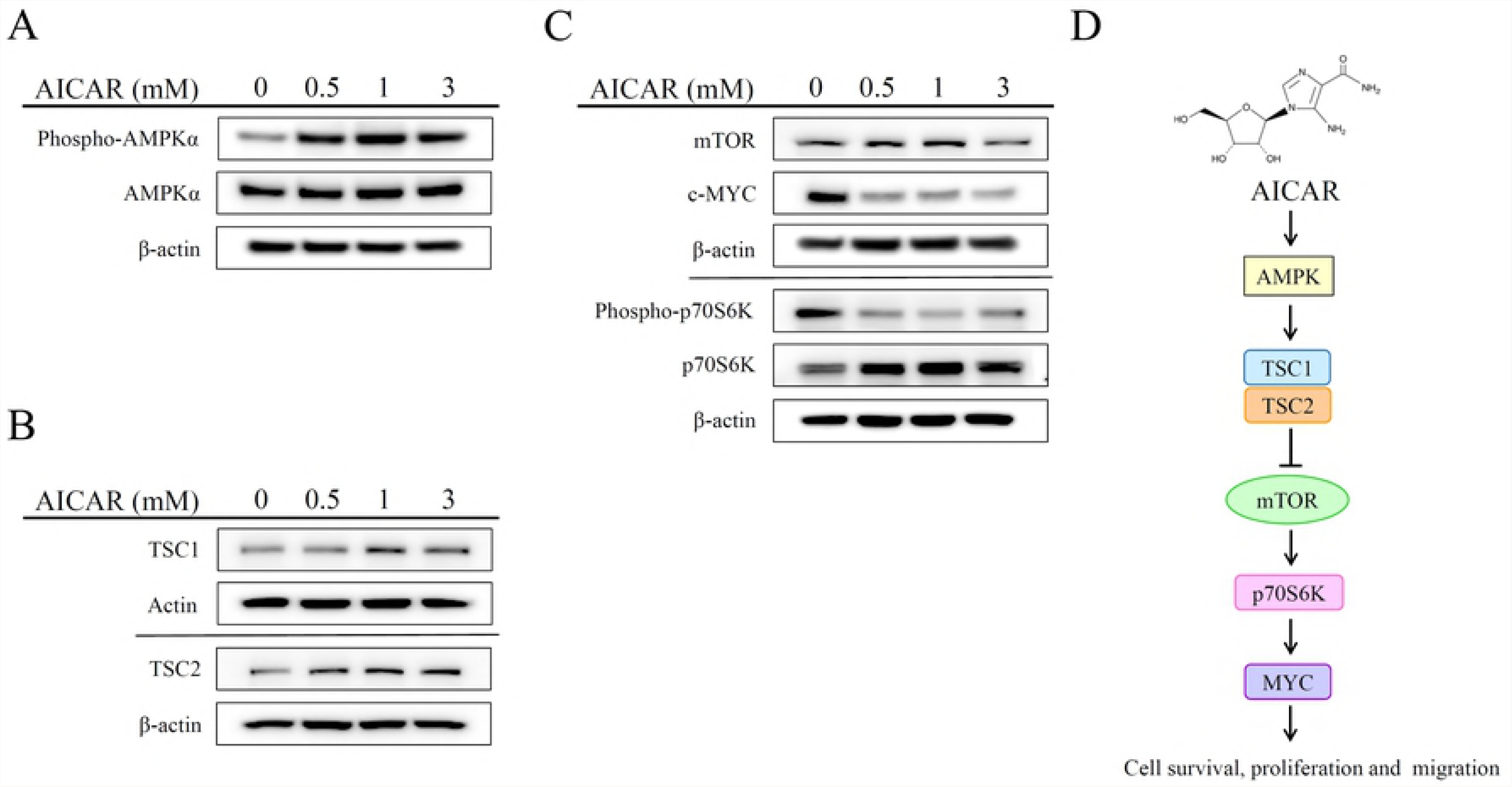
Effect of AICAR on the AMPK/mTOR-dependent pathway in 22Rv1 prostate cancer cells. Cells were treated with different concentrations of AICAR for 2 hr. The expression of (A) phospho-AMPK, AMPK, (B) TSC1 and TSC2 was examined by Western blot. Cells were treated with different concentrations of AICAR for 6 hr. (C) The expression of mTOR, cMYC, phosphor-p70S6K and p70S6K was determined by Western blot. Actin was used as a loading control in Western blot. The Western blotting results are representative of results obtained in three separate experiments. (D) Proposed mechanism of the anticancer effects induced by AICAR in 22Rv1 prostate cancer cells.

## Discussion

AMPK is an important regulator of cellular energy homeostasis [29]. Several agents have been demonstrated to activate AMPK, including metformin and phenformin, which increase the AMP:ATP ratio, the nucleoside AICAR which is metabolized to an AMP mimetic, and A769662, which is a direct activator of AMPK [30]. Although a number of studies indicated an anti-cancer role for AMPK [9, 11, 12, 14, 15, 31], the effect of an AMPK activator, AICAR, in regulating the pathological processes of prostate cancer remains enigmatic. Previous studies have shown that AICAR could exert cytotoxic effect on cancer cells via AMPK-dependent and/or AMPK-independent mechanisms [16, 31-33]. However, the cytotoxic effect on prostate cancer remains controversial. Guo et al. indicated that AICAR induces programmed necrosis, but not apoptosis, in prostate cancer cells, LNCaP, PC-3 and PC-82 cells through AMPK-independent pathways [21]. In contrast, Sauer et al. demonstrated that AICAR induces apoptosis of DU-145 prostate cancer cells through the AMPK/mTOR-dependent signaling pathway [16]. Our experimental results indicated that AICAR induces apoptosis, but not necrosis, in 22Rv1 prostate cancer cells. These results point out that AICAR inhibits cell growth and induces cell death in different prostate cancer cell lines probably through different mechanisms. In addition, AMPK is an upstream regulator in modulating the activation of TSC1 and TSC2, which consequently inhibits cell survival and proliferation via repressed mTOR-p70S6K-MYC signaling pathway [27, 28]. Our experimental results showed that AICAR enhances the expression of TSC1 and TSC2, whereas it reduces the expression of mTOR and MYC as well as decreases the phosphorylation of p70S6K in 22Rv1 prostate cancer cells. We suggest that AICAR induces apoptosis and inhibits migration of prostate cancer cells through an AMPK/mTOR-dependent pathway.

Cancer metastasis is the major cause of morbidity and mortality; approximately 80% of the men who have expired from prostate cancer possessed bone metastases [34]. Our experimental results indicated that AICAR suppresses TGF-β-induced cell migration and inhibits TGF-β-induced EMT by decreasing the expression of a mesenchymal-related marker, N-cadherin and enhancing the expression of an epithelial-related marker, E-cadherin in 22Rv1 prostate cancer cells. A recent study also indicated that treatment of AICAR suppresses migration and invasion in PC3 and PC3M prostate cancer cells [15]. These results implied that AICAR might have potential in inhibiting the metastatic activity in prostate cancer.

Docetaxel is a first-line treatment for CRPC, which confers survival advantages of approximately 2 months for patients with low overall survival benefit [35]. However, treatment with docetaxel usually causes adverse side effects including hair loss, myelosuppression, neurotoxicity, diarrhea etc. Our experimental results showed that AICAR has synergistic effect with docetaxel treatment. These results suggest that AICAR increases the sensitization of prostate cancer cells to docetaxel-induced apoptosis, which might have benefit for reducing toxicity of chemotherapy in prostate cancer patients.

In summary, our experimental results indicated that AICAR inhibits cell growth, induces apoptosis, attenuates cell migration, enhances chemosensitivity to docetaxel, and suppresses the activation of the AMPK/mTOR-dependent signaling pathway. These results suggest that AICAR appears as a new potential anticancer agent for treating prostate cancer.

## Author Contributions

**Conceptualization: CS** SF WC MH CL.

**Data curation:**CS KLH SW HY SH PL KHH FC IL.

**Formal analysis:** CS KLH SW FC.

**Funding acquisition:** KLH WC MH CL.

**Investigation:** CS KLH MH CL.

**Methodology:** SH MH CL.

**Project administration:** MH CL.

**Resources:** MH CL.

**Supervision:** MH CL.

**Validation:** CS KLH FC.

**Visualization:** CS KLH MH CL.

**Writing – original draft:** CS CL.

**Writing – review & editing:** HY SF SH MH CL

## References

1. Siegel RL, Miller KD, Jemal A. Cancer statistics, 2018. CA Cancer J Clin. 2018;68(1):7–30. doi: 10.3322/caac.21442. PubMed PMID: 29313949.

2. Litwin MS, Tan HJ. The Diagnosis and Treatment of Prostate Cancer: A Review. JAMA. 2017;317(24):2532–42. doi: 10.1001/jama.2017.7248. PubMed PMID: 28655021.

3. Marshall S, Taneja S. Focal therapy for prostate cancer: The current status. Prostate Int. 2015;3(2):35–41. doi: 10.1016/j.prnil.2015.03.007. PubMed PMID: 26157765; PubMed Central PMCID: PMCPMC4494637.

4. Asmane I, Ceraline J, Duclos B, Rob L, Litique V, Barthelemy P, et al. New strategies for medical management of castration-resistant prostate cancer. Oncology. 2011;80(1-2):1–11. doi: 10.1159/000323495. PubMed PMID: 21577012.

5. Grahame Hardie D. Regulation of AMP-activated protein kinase by natural and synthetic activators. Acta Pharm Sin B. 2016;6(1):1–19. doi: 10.1016/j.apsb.2015.06.002. PubMed PMID: 26904394; PubMed Central PMCID: PMCPMC4724661.

6. Bai P, Nagy L, Fodor T, Liaudet L, Pacher P. Poly(ADP-ribose) polymerases as modulators of mitochondrial activity. Trends Endocrinol Metab. 2015;26(2):75–83. doi: 10.1016/j.tem.2014.11.003. PubMed PMID: 25497347.

7. Hardie DG. AMP-activated/SNF1 protein kinases: conserved guardians of cellular energy. Nat Rev Mol Cell Biol. 2007;8(10):774–85. doi: 10.1038/nrm2249. PubMed PMID: 17712357.

8. Hsu PP, Sabatini DM. Cancer cell metabolism: Warburg and beyond. Cell. 2008;134(5):703–7. doi: 10.1016/j.cell.2008.08.021. PubMed PMID: 18775299.

9. Icard P, Lincet H. A global view of the biochemical pathways involved in the regulation of the metabolism of cancer cells. Biochim Biophys Acta. 2012;1826(2):423–33. doi: 10.1016/j.bbcan.2012.07.001. PubMed PMID: 22841746.

10. Kroemer G. Mitochondria in cancer. Oncogene. 2006;25(34):4630–2. doi: 10.1038/sj.onc.1209589. PubMed PMID: 16892077.

11. Kishton RJ, Barnes CE, Nichols AG, Cohen S, Gerriets VA, Siska PJ, et al. AMPK Is Essential to Balance Glycolysis and Mitochondrial Metabolism to Control T-ALL Cell Stress and Survival. Cell Metab. 2016;23(4):649–62. doi: 10.1016/j.cmet.2016.03.008. PubMed PMID: 27076078; PubMed Central PMCID: PMCPMC4832577.

12. Fodor T, Szanto M, Abdul-Rahman O, Nagy L, Der A, Kiss B, et al. Combined Treatment of MCF-7 Cells with AICAR and Methotrexate, Arrests Cell Cycle and Reverses Warburg Metabolism through AMP-Activated Protein Kinase (AMPK) and FOXO1. PLoS One. 2016;11(2):e0150232. doi: 10.1371/journal.pone.0150232. PubMed PMID: 26919657; PubMed Central PMCID: PMCPMC4769015.

13. Cheng X, Kim JY, Ghafoory S, Duvaci T, Rafiee R, Theobald J, et al. Methylisoindigo preferentially kills cancer stem cells by interfering cell metabolism via inhibition of LKB1 and activation of AMPK in PDACs. Mol Oncol. 2016. doi: 10.1016/j.molonc.2016.01.008. PubMed PMID: 26887594.

14. Park SY, Lee YK, Kim HJ, Park OJ, Kim YM. AMPK interacts with beta-catenin in the regulation of hepatocellular carcinoma cell proliferation and survival with selenium treatment. Oncol Rep. 2016;35(3):1566–72. doi: 10.3892/or.2015.4519. PubMed PMID: 26707164.

15. Choudhury Y, Yang Z, Ahmad I, Nixon C, Salt IP, Leung HY. AMP-activated protein kinase (AMPK) as a potential therapeutic target independent of PI3K/Akt signaling in prostate cancer. Oncoscience. 2014;1(6):446–56. PubMed PMID: 25594043; PubMed Central PMCID: PMCPMC4284621.

16. Sauer H, Engel S, Milosevic N, Sharifpanah F, Wartenberg M. Activation of AMP-kinase by AICAR induces apoptosis of DU-145 prostate cancer cells through generation of reactive oxygen species and activation of c-Jun N-terminal kinase. Int J Oncol. 2012;40(2):501–8. doi: 10.3892/ijo.2011.1230. PubMed PMID: 22002081.

17. Fogarty S, Hardie DG. Development of protein kinase activators: AMPK as a target in metabolic disorders and cancer. Biochim Biophys Acta. 2010;1804(3):581–91. doi: 10.1016/j.bbapap.2009.09.012. PubMed PMID: 19778642.

18. Jacobs RL, Lingrell S, Dyck JR, Vance DE. Inhibition of hepatic phosphatidylcholine synthesis by 5-aminoimidazole-4-carboxamide-1-beta-4-ribofuranoside is independent of AMP-activated protein kinase activation. J Biol Chem. 2007;282(7):4516–23. doi: 10.1074/jbc.M605702200. PubMed PMID: 17179149.

19. Lopez JM, Santidrian AF, Campas C, Gil J. 5-Aminoimidazole-4-carboxamide riboside induces apoptosis in Jurkat cells, but the AMP-activated protein kinase is not involved. Biochem J. 2003;370(Pt 3):1027–32. doi: 10.1042/BJ20021053. PubMed PMID: 12452797; PubMed Central PMCID: PMCPMC1223217.

20. Xiang X, Saha AK, Wen R, Ruderman NB, Luo Z. AMP-activated protein kinase activators can inhibit the growth of prostate cancer cells by multiple mechanisms. Biochem Biophys Res Commun. 2004;321(1):161–7. doi: 10.1016/j.bbrc.2004.06.133. PubMed PMID: 15358229.

21. Guo F, Liu SQ, Gao XH, Zhang LY. AICAR induces AMPK-independent programmed necrosis in prostate cancer cells. Biochem Biophys Res Commun. 2016. doi: 10.1016/j.bbrc.2016.04.077. PubMed PMID: 27103440.

22. Duan F, Li Y, Chen L, Zhou X, Chen J, Chen H, et al. Sulfur inhibits the growth of androgen-independent prostate cancer in vivo. Oncol Lett. 2015;9(1):437–41. doi: 10.3892/ol.2014.2700. PubMed PMID: 25436005; PubMed Central PMCID: PMCPMC4247018.

23. Boulares AH, Yakovlev AG, Ivanova V, Stoica BA, Wang G, Iyer S, et al. Role of poly(ADP-ribose) polymerase (PARP) cleavage in apoptosis. Caspase 3-resistant PARP mutant increases rates of apoptosis in transfected cells. J Biol Chem. 1999;274(33):22932–40. PubMed PMID: 10438458.

24. Barrett CS, Millena AC, Khan SA. TGF-beta Effects on Prostate Cancer Cell Migration and Invasion Require FosB. Prostate. 2017;77(1):72–81. doi: 10.1002/pros.23250. PubMed PMID: 27604827; PubMed Central PMCID: PMCPMC5286811.

25. Petrylak DP, Tangen CM, Hussain MH, Lara PN, Jr., Jones JA, Taplin ME, et al. Docetaxel and estramustine compared with mitoxantrone and prednisone for advanced refractory prostate cancer. N Engl J Med. 2004;351(15):1513–20. doi: 10.1056/NEJMoa041318. PubMed PMID: 15470214.

26. Tannock IF, de Wit R, Berry WR, Horti J, Pluzanska A, Chi KN, et al. Docetaxel plus prednisone or mitoxantrone plus prednisone for advanced prostate cancer. N Engl J Med. 2004;351(15):1502–12. doi: 10.1056/NEJMoa040720. PubMed PMID: 15470213.

27. Williams T, Courchet J, Viollet B, Brenman JE, Polleux F. AMP-activated protein kinase (AMPK) activity is not required for neuronal development but regulates axogenesis during metabolic stress. Proc Natl Acad Sci U S A. 2011;108(14):5849–54. doi: 10.1073/pnas.1013660108. PubMed PMID: 21436046; PubMed Central PMCID: PMCPMC3078367.

28. Zhou W, Marcus AI, Vertino PM. Dysregulation of mTOR activity through LKB1 inactivation. Chin J Cancer. 2013;32(8):427–33. doi: 10.5732/cjc.013.10086. PubMed PMID: 23668926; PubMed Central PMCID: PMCPMC3845579.

29. Khan AS, Frigo DE. A spatiotemporal hypothesis for the regulation, role, and targeting of AMPK in prostate cancer. Nat Rev Urol. 2017;14(3):164–80. doi: 10.1038/nrurol.2016.272. PubMed PMID: 28169991; PubMed Central PMCID: PMCPMC5672799.

30. Hawley SA, Ross FA, Chevtzoff C, Green KA, Evans A, Fogarty S, et al. Use of cells expressing gamma subunit variants to identify diverse mechanisms of AMPK activation. Cell Metab. 2010;11(6):554–65. doi: 10.1016/j.cmet.2010.04.001. PubMed PMID: 20519126; PubMed Central PMCID: PMCPMC2935965.

31. Chen L, Han F, Qu H, Yan H, Ren L, Yang S. Combination therapy with 5-amino-4-imidazolecarboxamide riboside and arsenic trioxide in acute myeloid leukemia cells involving AMPK/TSC2/mTOR pathway. Pharmazie. 2013;68(2):117–23. PubMed PMID: 23469683.

32. Santidrian AF, Gonzalez-Girones DM, Iglesias-Serret D, Coll-Mulet L, Cosialls AM, de Frias M, et al. AICAR induces apoptosis independently of AMPK and p53 through up-regulation of the BH3-only proteins BIM and NOXA in chronic lymphocytic leukemia cells. Blood. 2010;116(16):3023–32. doi: 10.1182/blood-2010-05-283960. PubMed PMID: 20664053.

33. Su RY, Chao Y, Chen TY, Huang DY, Lin WW. 5-Aminoimidazole-4-carboxamide riboside sensitizes TRAIL- and TNF{alpha}-induced cytotoxicity in colon cancer cells through AMP-activated protein kinase signaling. Mol Cancer Ther. 2007;6(5):1562–71. doi: 10.1158/1535-7163.MCT-06-0800. PubMed PMID: 17513605.

34. Bubendorf L, Schopfer A, Wagner U, Sauter G, Moch H, Willi N, et al. Metastatic patterns of prostate cancer: an autopsy study of 1,589 patients. Hum Pathol. 2000;31(5):578–83. PubMed PMID: 10836297.

35. Poorthuis MHF, Vernooij RWM, van Moorselaar RJA, de Reijke TM. First-line non-cytotoxic therapy in chemotherapy-naive patients with metastatic castration-resistant prostate cancer: a systematic review of 10 randomised clinical trials. BJU Int. 2017;119(6):831–45. doi: 10.1111/bju.13764. PubMed PMID: 28063195.

